# Strange effects of activation dynamics on musculoskeletal trajectory optimization

**DOI:** 10.1101/2025.01.30.635759

**Authors:** Antonie J. (Ton) van den Bogert

**Affiliations:** Cleveland State University

## Abstract

Muscle activation dynamics is usually described with a nonlinear differential equation, such that the activation occurs faster than deactivation. When a muscle excitation input switches rapidly between two values, this produces a “pumping” effect that raises the activation above the mean excitation input. This effect will favor rapid switching strategies when excitation is used in the optimization objective for human movement. The optimal strategy has infinitely fast switching, and is difficult to obtain with direct collocation methods. This problem can be eliminated by using activation, rather than excitation, in the optimization objective.

## 1. Introduction

Predictive simulations of running can be performed by trajectory optimization on a musculoskeletal model, in which a weighted sum of squared tracking errors and muscle excitations is minimized (van den Bogert et al., 2012). In a recent project, we attempted to improve the accuracy of these optimizations by increasing the number of collocation points, as well as very tight tolerance setting for the solver. As the solver progressed to tighter tolerance, optimal excitation input for most muscles developed a rapid on/off switching strategy (Figure 1). As the optimal controls were becoming more and more discontinuous, the cost function slowly decreased.

**Figure 1.**
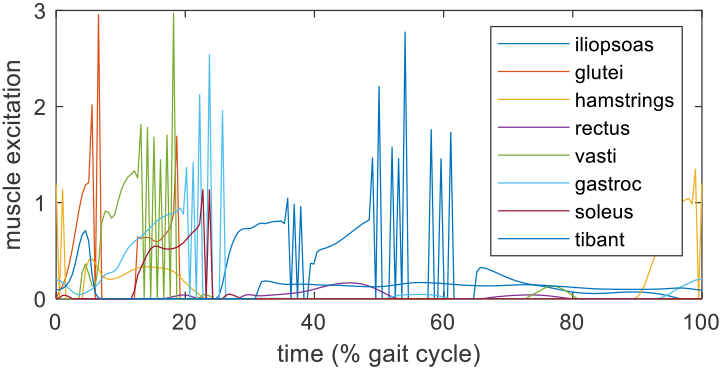
Optimal controls for a running simulation, using 200 collocation points and a backward Euler discretization.

It was suspected that this behavior was related to the muscle activation dynamics model.

### 2. The “pumping” effect of activation dynamics

Muscle activation dynamics is usually described with a nonlinear differential equation. For example, McLean et al. (2003) used:

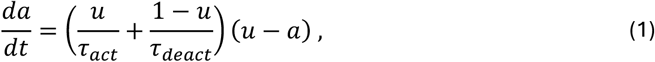

where *u* is the excitation input, *a* is the activation state, and *τ* are the time constants for activation and deactivation. The activation model of De Groote et al. (2016) is commonly used and probably more accurate. All models for activation dynamics have an activation time constant parameter that is smaller (faster) than the deactivation time constant. This creates an interesting “rectification” or “pumping” effect when the input is switched rapidly between two values. This increases the mean activation to a level larger than the mean excitation (Figure 2).

**Figure 2.**
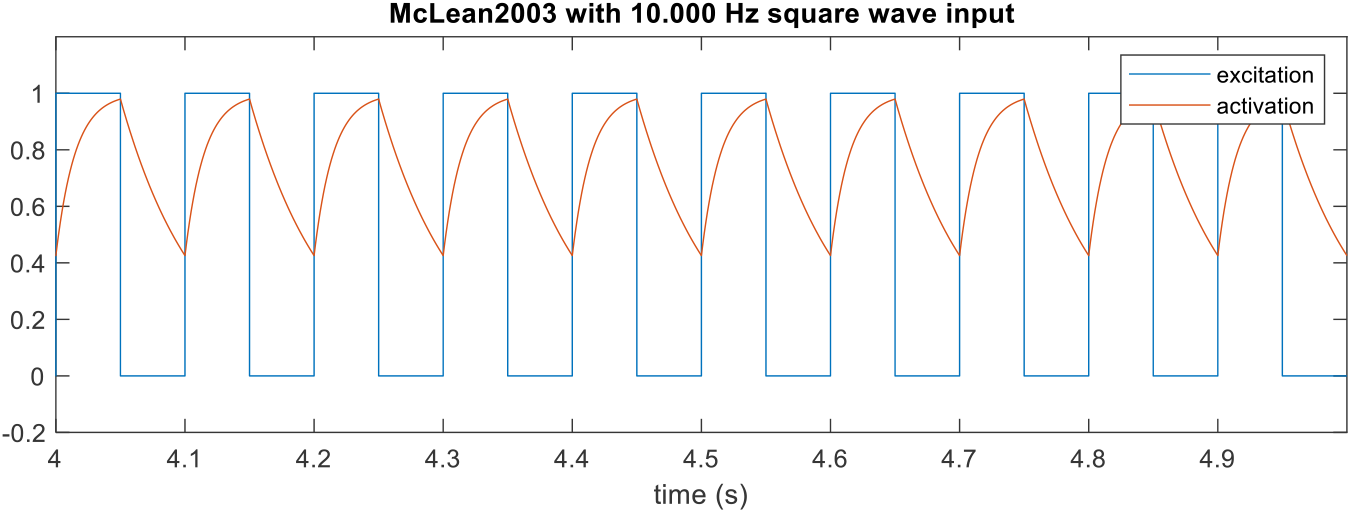
Simulated activation response with the McLean et al. (2003) model, and activation/deactivation time constants of 15 and 60 ms, respectively.

To further explore this phenomenon, we simulated the activation models of McLean et al. (2003) and De Groote et al., (2016) with square wave inputs of several frequencies. Time constants of 15 ms and 60 ms were used in both models. Square wave input signals always have the same cost function value, regardless of frequency, when cost is any time-integrated function of excitation.

In De Groote’s model, the tanh steepness parameter (*b*) was changed from 0.1 to 10.0, otherwise the activation and deactivation speeds did not differ much. I suspect that this was the original intention of the model.

The results show the expected effect, that higher frequencies produce higher mean activations (Figures 3-4).

**Figure 3.**
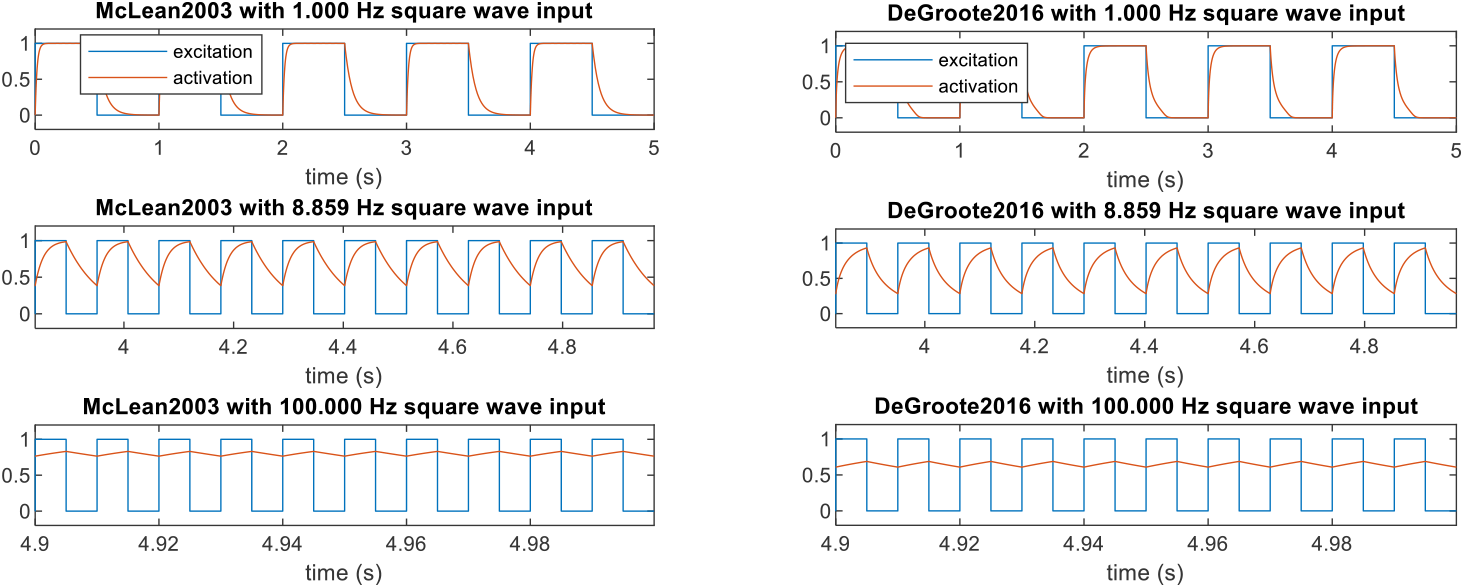
Activation responses from two activation dynamics models, driven by square wave excitations.

### 2. Analytical optimal control solution

The simulations presented in the previous section suggest that an optimal excitation signal will always have infinite switching frequency.

We consider the problem of producing a mean muscle activation of 0.5, with minimal control effort. Effort is defined as the mean of squared excitation. The model of McLean et al. (2003) was used.

We parameterize the bang-bang excitation signal by frequency (*f*) and duty cycle (*c*).

When the frequency is infinitely large, the activation output (*a*) will be constant because it does not have time to rise and fall more than an infinitesimal amount.

During the infinitesimally short activation time (*c*/*f*), the activation will rise by the activation rate (*da*/*dt*), multiplied by the duration (*c*/*f*):

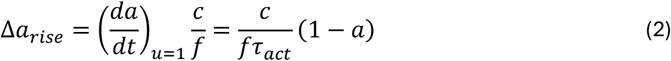

Similarly, during the deactivation time ((1 − *c*)/*f*), the activation will fall by this amount:

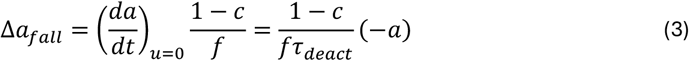

Steady state is achieved when the amounts of rise and fall are equal and opposite. This produces a relationship between the steady state output (*a*) and the duty cycle (*c*):

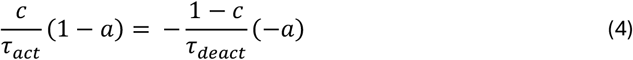

We solve for *c*, i.e. the duty cycle required to achieve a constant activation *a*:

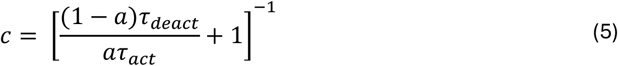

For *τ*_*act*_ = 15 ms and *τ*_*deact*_ = 60 ms, the required duty cycle to achieve an activation *a*=0.5 will be exactly 0.2.

An analytical solution for the more complex model of De Groote et al. (2016) can be derived similarly.

In all cases, the mean squared excitation cost will be exactly equal to the duty cycle:

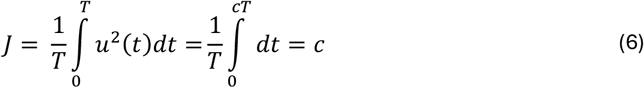

It is expected that a numerical trajectory optimization method will have difficulty approximating such a solution where the control input switches infinitely fast between two values.

## 3. Numerical optimal control solution

Direct collocation was used to find the optimal excitation input to achieve a mean activation of 0.5.

The problem was formulated as: minimize:

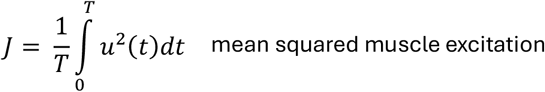

subject to:

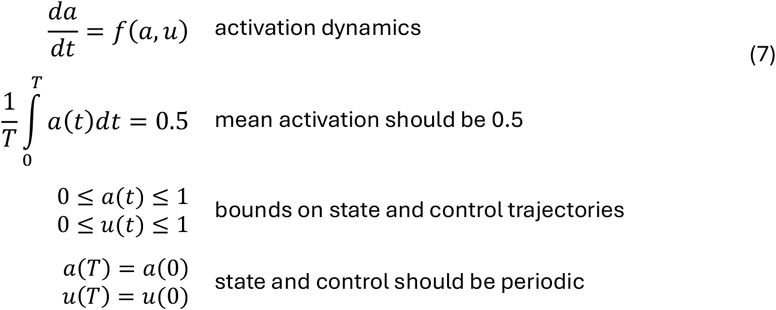

where *f*() is the activation dynamics model of McLean et al. (2003) with time constants of 15 and 60 ms. Simulation time (*T*) was 1 second.

The problem was time-discretized into 200 equally spaced collocation points, and the trapezoidal formula was used to approximate the differential equation. The IPOPT solver was used with a random initial guess and default tolerances.

IPOPT solved the problem in 5865 iterations (taking 17 seconds). The solution (Figure 5, left) showed a switching strategy, but was obviously not able to switch infinitely fast, or achieve the optimal duty cycle, due to the limited resolution of the temporal mesh. The number of iterations indicates that the problem is hard to solve. The timing of the switching was irregular, which would not be expected in a true optimal strategy. The cost function value in the numerical solution was 0.2030, slightly larger than the analytical optimum of 0.2. When the Backward Euler discretization was used, it took longer to solve (12760 iterations), the cost function was worse (0.2215), and the solution still showed the switching strategy.

The analytical solution, with duty cycle 0.2 (and cost function value 0.2) was then simulated with a frequency of 1000 Hz, generating a mean activation of 0.499996 (Figure 5, right). Achieving a mean activation of exactly 0.5 would require an infinite switching frequency.

The optimization was repeated with 1000 collocation points. The solution was obtained in 54352 iterations (taking 968 seconds) and achieved a cost function value of 0.2009. With the increased temporal resolution, the solution came closer to the analytical optimum but at great cost in computation time.

## 4. Discussion

When muscle excitation is used in a cost function for musculoskeletal control, the true optimal control solution has infinitely fast switching of the control input. The numerical solution will try to approximate this solution, resulting in a suboptimal solution with “spiky” controls, and excessive computation time when accuracy is important.

This behavior has not been reported before, and there are several possible reasons.

First, temporal meshes are typically 50-100 points for a gait cycle, i.e. a time step of 10-20 ms between collocation points. When the time step is comparable to the time constants, the activation dynamics is only very crudely simulated. We first noticed the rapid switching for running at 4.3 m/s, which has a shorter gait cycle, using 200 collocation points, resulting in about 3.4 ms between collocation points. This time step is small enough that the activation dynamics is accurately simulated and therefore matters more. To test this idea, the “mean activation = 0.5” problem was solved with 50 collocation points per second. With the trapezoidal discretization, the solution was still “spiky”. However, with the backward Euler method, which we often used in the past, the solution was a constant u(t)=0.5 and found very quickly in 16 iterations. Discretization formulas that are asymmetric in time, such as Backward Euler (Nitschke et al., 2020), 3^rd^ order Radau (Falisse et al., 2019), or Hermite-Simpson (Dembia et al., 2019) can introduce numerical damping which cancels the favorability of rapid switching solutions, unless the time step is sufficiently small.

Second, musculoskeletal optimal control often uses an additional “regularization” cost term which penalizes rapid variations in states and/or controls (Falisse et al., 2019; Nitschke et al. 2020). This penalization is generally justified by the idea that rapid variations don’t matter for performance, and by weighting this term such that it is extremely small in the final optimal movement. It is thought that this term only helps to find the optimum, and does not actually change the optimum. For this reason, this term is often not even mentioned in the Methods section of a publication. However, in this report we showed that penalizing of rapid switching can cause the true optimum, which has the switching strategy, to be missed.

Third, this problem would not have been noticed in work where activation (rather than excitation) is used to define the effort cost (e.g. Ackermann and van den Bogert, 2010). In such work, the activation state trajectory is really what is optimized, and the activation dynamics are only there to ensure that the activation state trajectories are feasible, i.e. can be achieved with excitations that are bounded between zero and one. I believe that this is the correct approach. When activation cost was used in the simulations of running (see Introduction), the optimal controls were more smooth and the solution process appeared to be faster.

Fourth, the problem only occurs when there is substantial difference between activation and deactivation speeds. The model of De Groote et al. (2016) is currently the most popular model for optimal control of human movement. However, as published and as currently implemented in OpenSim (DeGrooteFregly2016Muscle), this model has almost no difference between activation and deactivation speed, eliminating the benefit of excitation switching (Figure 6). The issue shown in this report might not appear when using the OpenSim implementation.

McLean et al. (2003) has activation/deactivation rate depending on the input *u*, while it should depend on the difference between *u* and *a*, as in Groote et al., (2016). However, both have the “pumping” effect which is a real property of muscles that should be represented in models.

The McLean2003 model also has a problem when the excitation is allowed to go above 1, as is sometimes done when muscle strength, derived from cadaver measurements, is not sufficient to perform a high intensity task. Equation (1) shows that this would make the time constant shorter than *τ*_*act*_, which is not physiological. We propose the following improvement of the McLean2003 model:

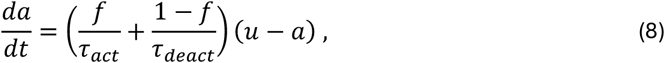

where f is smooth sigmoid function *f*(*u* − a) that is approximately 1 for activation (*u* > a) and 0 for deactivation (*u* < a). De Groote et al. (2016) used a logistic function. We propose an algebraic function, which does not saturate as quickly, which should be advantageous for numerical solution methods (Figure 7):

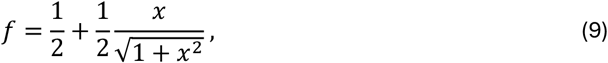

where *x* = b(*u* − a), with *b* = 10.

The square wave response of the activation model from equation (8) is plotted in Figure 6, along with the other models.

This improved version of the McLean et al (2003) activation model still produces rapid switching solutions in optimal control problems. Surprisingly, the collocation method solves much more quickly than the original problem. The local optimum suggests that the optimal control still wants to switch rapidly, but no longer between 0 and 1 (Figure 8). It may be that IPOPT solves this problem more easily because the control signal does not approach the bounds. A square wave solution was found by shooting. It rapidly switched between 0.1998 and 0.8002 with a duty cycle of 0.2002 (Figure 8).

The DeGroote2016 model had a similar optimal control solution, with the control signal rapidly switching, but not hitting the lower and upper bounds. Table 1 summarizes the results for all models.

**Table 1.** Parameters of the optimal square wave control signal for producing a mean activation of 0.5.

| Model | $u_{\min}$ | $u_{\max}$ | duty cycle |
| --- | --- | --- | --- |
| McLean2003 | 0.0000 | 1.0000 | 0.2000 |
| McLean2003improved | 0.1998 | 0.8002 | 0.2002 |
| DeGroote2016 (b=10) | 0.2662 | 0.7338 | 0.2850 |

We conclude that, for all activation models, the simple problem of generating a specified mean activation (or mean muscle force) had a solution with infinitely fast switching of the control input.

## 5. Recommendation

Apart from non-smooth optimal control solutions, which do not look nice or convincing, we found that numerical methods have trouble finding these solutions. Therefore, it would be best to avoid the issue.

Based on the findings in this report, **it is recommended that muscle excitations are not used in cost functions for optimal control of human movement. Instead, activation should be used**. There is also a good physiological justification for this: activation determines energy consumption and fatigue, which are likely to be factors that humans consider when optimizing movement.

Excitation, on the other hand, is an electrical neural signal that does not cost anything, and may not be relevant in optimal control of human movement. A regularization term might still be beneficial to encourage smoothness and uniqueness in the optimal controls, but would no longer alter the solution fundamentally.

This recommendation may also extend to engineering applications, for systems with nonlinear dynamics. In linear systems, a quadratic control cost will always favor smooth controls. However, we have shown here that this is not generally the case for nonlinear systems. Model Predictive Control relies on performing trajectory optimizations in real time, and could benefit from avoiding cost functions that favor solutions which are inaccurate or difficult to find with numerical methods.

## MATLAB code

The Matlab code can be downloaded from https://github.com/csu-hmc/activation-dynamics.

The results presented in this report were produced with Matlab version 2024a, but other versions should work equally well.

To solve the optimal control problem numerically (Figure 5), either IPOPT or the Matlab Optimization Toolbox (fmincon) must be installed. If IPOPT is not installed, fmincon will be used. Instructions:

>> actsim % produces the result presented in Figure 2.

>> actdyntest % produces the results presented in Figures 3 and 4.

**Figure 4.**
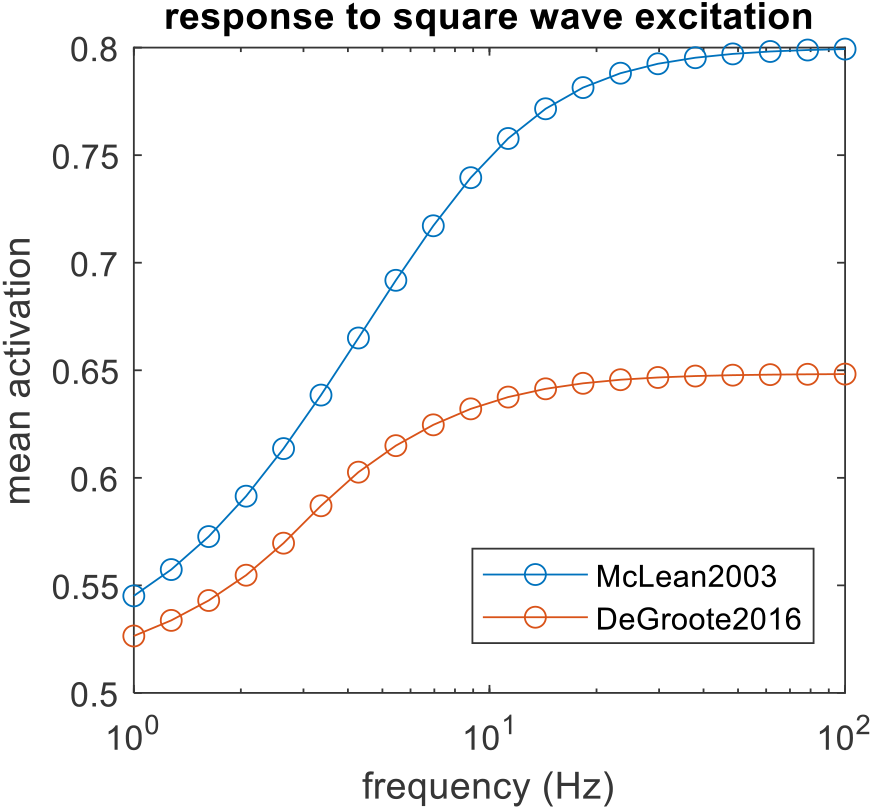
Mean activation from two activation dynamics models, driven by square wave excitations.

>> optimize % produces the results presented in Figure 5.

**Figure 5.**
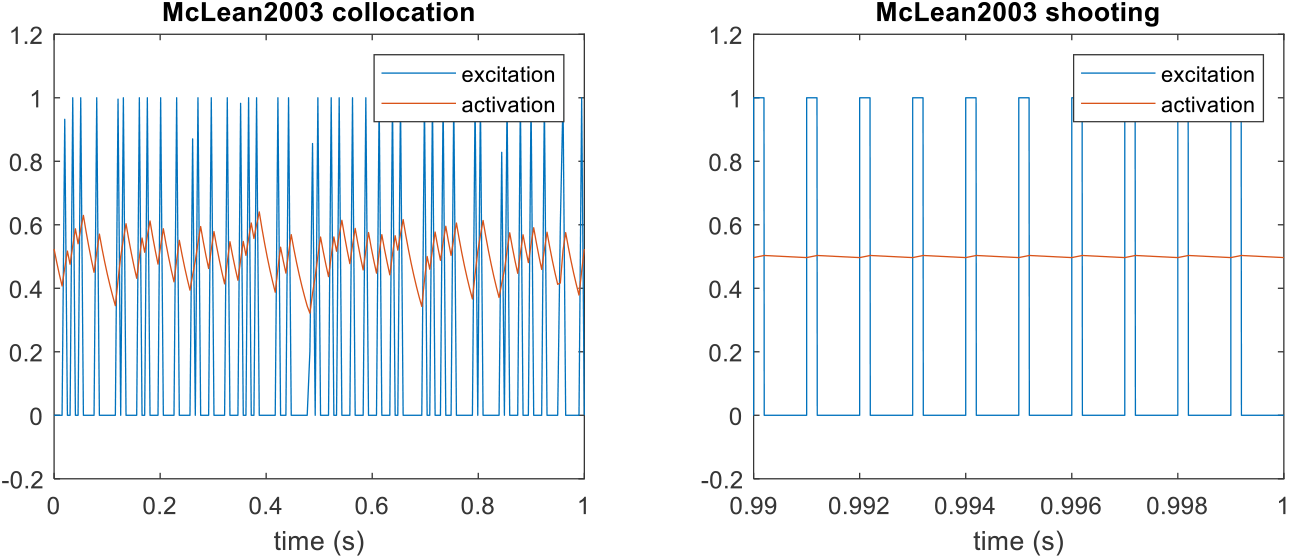
Numerical optimal control solution for the problem of generating a mean activation of 0.5 with the model of McLean et al. (2003). The analytical solution for a 1000 Hz switching frequency is shown on the right. Note the different time scales on the horizontal axis.

**Figure 6.**
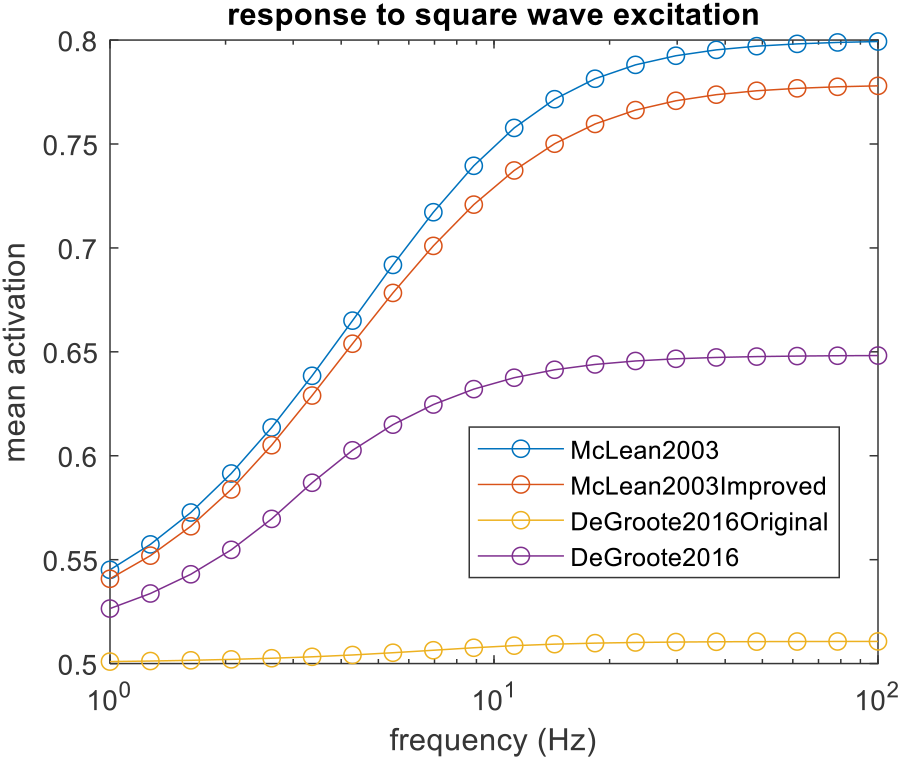
Mean activation generated by square wave excitation in four activation models. The original version of De Groote et al. (2016) has the parameter value *b*=0.1. The version used earlier in this report has *b*=10, which we suspect is the intended value. The improved version of McLean2003 has an activation rate that depends on u-a, rather than u, and does not result in unstable dynamics when u goes above 1.

**Figure 7.**
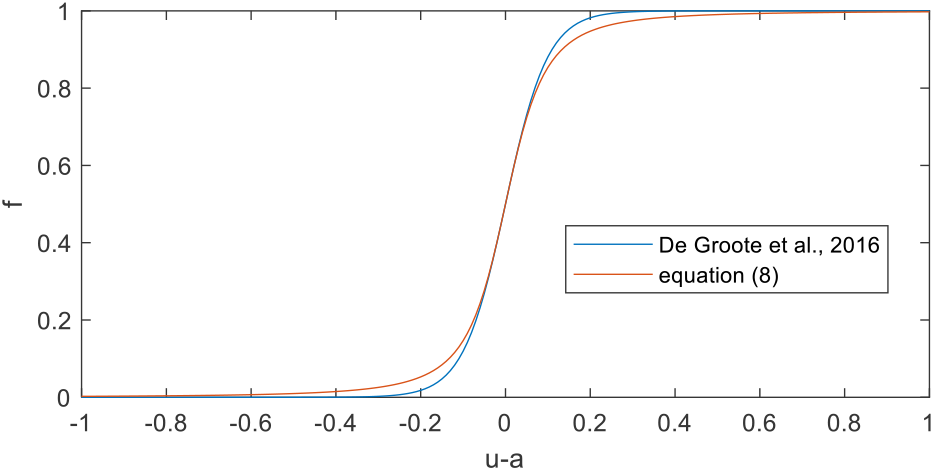
Sigmoid functions for switching between activation (f=1) and deactivation (f=0).

**Figure 8.**
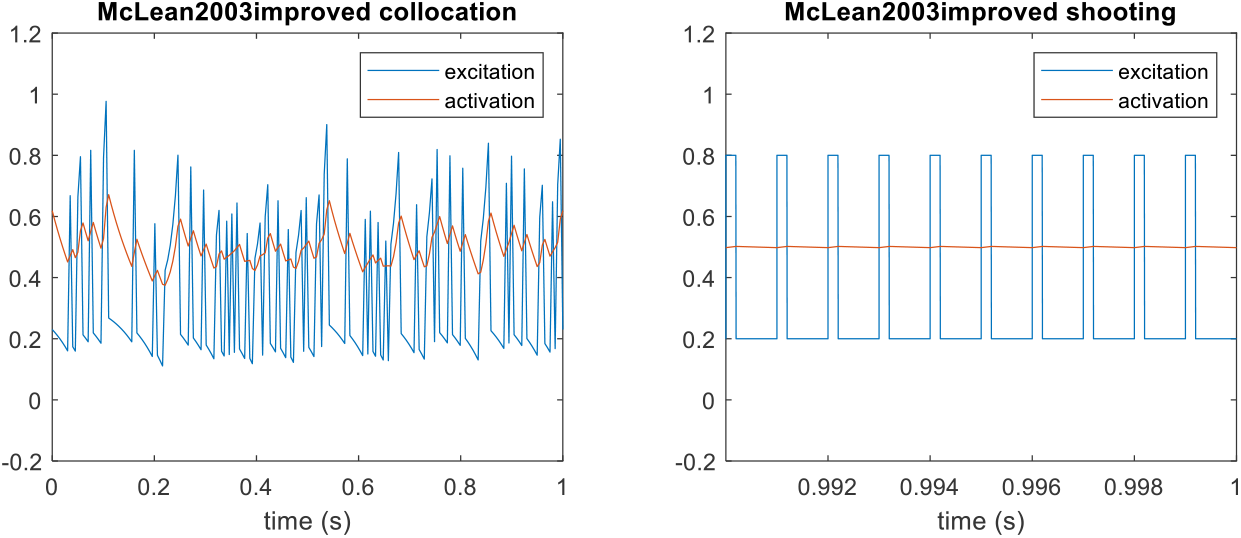
Optimal controls for producing a mean activation of 0.5 with the activation dynamics model of equation (8). The collocation solution (left) has a mean-squared-controls cost of 0.1686, while the square wave solution (right) has a cost of 0.1600.

## Notes

https://github.com/csu-hmc/activation-dynamics

### Competing Interest Statement

The authors have declared no competing interest.

### Summary of Updates

A proposed new activation model was added, with corresponding results of simulation and analysis.

